# Unicellular associative conditioning: an interspecies analysis

**DOI:** 10.1101/2020.10.19.346007

**Authors:** Jose Carrasco-Pujante, Carlos Bringas, Iker Malaina, Maria Fedetz, Luis Martínez, Gorka Pérez-Yarza, María Dolores Boyano, Mariia Berdieva, Andrew Goodkov, José I. López, Shira Knafo, Ildefonso M. De la Fuente

**Affiliations:** Department of Physiology and Cell Biology, Faculty of Health Sciences, and The National Institute for Biotechnology in the Negev, Ben-Gurion University of the Negev, Beer-Sheva 8410501, Israel; Department of Cell Biology and Histology, Faculty of Medicine and Nursing, University of the Basque Country, UPV/EHU, Leioa 48940, Spain; Department of Mathematics, Faculty of Science and Technology, University of the Basque Country, UPV/EHU, Leioa 48940, Spain; Department of Cell Biology and Immunology, Institute of Parasitology and Biomedicine “López-Neyra”, CSIC, Granada 18016, Spain; Basque Center of Applied Mathematics (BCAM) Bilbao 48009, Spain; Laboratory of Cytology of Unicellular Organisms, Institute of Cytology Russian Academy of Science St. Petersburg 194064, Russia; Department of Pathology, Cruces University Hospital, Biocruces-Bizkaia Health Research Institute, Barakaldo 48903, Spain; Biophysics Institute, CSIC-UPV/EHU, Campus, University of the Basque Country (UPV/EHU), Leioa 48940, Spain; Ikerbasque, Basque Foundation for Science, Bilbao 48013, Spain; Department of Nutrition, CEBAS-CSIC Institute, Espinardo University Campus, Murcia 30100, Spain

**Author notes:** **Corresponding Authors**: Correspondence and requests for materials should be addressed to Shira Knafo or Ildefonso M. de la Fuente. **Ildefonso M. de la Fuente** Instituto CEBAS-CSIC. Campus Universitario de Espinardo. Espinardo. 30100 Murcia. ESPAÑA. **Shira Knafo** Department of Physiology and Cell Biology, Faculty of Health Sciences, and The National Institute for Biotechnology in the Negev, Ben-Gurion University of the Negev, Beer-Sheva 8410501, Israel. **Equal contribution** (Jose Carrasco-Pujante, Carlos Bringas).

## Abstract

The capacity to learn new systemic behaviour is a fundamental issue to understand the adaptive mechanisms involved in cellular evolution. We have recently observed, in a preliminary analysis, the emergence of conditioned behaviour in individual amoebae cells. In these experiments, cells were able to acquire new migratory conduct and remember it for long periods of their cellular cycle, forgetting it later on. Here, following a similar conceptual framework of Pavlov’s experiments, we have exhaustively studied the migration trajectories of more than 2000 individual cells belonging to three different species: *Amoeba proteus, Metamoeba leningradensis*, and *Amoeba borokensis*. Fundamentally, we have analysed several properties of conditioned cells, such as the intensity of the responses, the directionality persistence, the total distance traveled, the directionality ratio, the average speed, and the persistence times. We have observed that these three species can modify the systemic response to a specific stimulus by associative conditioning. Our main analysis shows that such new behaviour is very robust and presents a similar structure of migration patterns in the three species, which was characterized by the presence of conditioning for long periods, remarkable straightness in their trajectories and strong directional persistence. Our quantitative results, compared with other studies on complex cellular responses in bacteria, protozoa, fungus-like organisms and metazoans, allow us to conclude that cellular associative conditioning might be a widespread characteristic of unicellular organisms. This finding could be essential to understand some key evolutionary principles involved in increasing the cellular adaptive fitness to microenvironments.

## Introduction

An essential question in evolutionary studies is to know if individual cells are capable of learning, thereby acquiring new adaptive systemic patterns to respond to changes in the external medium.

Associative learning and memory are fundamental cognitive properties of multicellular organisms endowed with nervous systems to acquire critical information for survival, developing new complex behaviour by the association of different stimuli and/or responses. This property is ubiquitous in many species, from cephalopods to humans^1^. Recently, we observed in a preliminary study the emergence of cellular conditioning in *Amoeba proteus* sp., which modified their migratory behaviour by an association of external stimuli acquiring a new systemic response^2^.

In continuation of this research and following a similar conceptual framework of Pavlov’s experiments, we have studied here the migration trajectories of more than 2000 individual cells belonging to three different species (*Amoeba proteus*, sp., *Metamoeba leningradensis*, sp. and *Amoeba borokensis*, sp.). First, we have analysed the locomotion movements under three external conditions: in absence of stimuli, in an electric field (galvanotaxis), and under chemotactic gradient (chemotaxis). Next, we have conditioned a great number of cells, applying simultaneously galvanotactic and chemotactic stimuli, and we verified that most of them (78%) can acquire a new systemic behaviour (movement to the anode when the habitual behaviour under galvanotactic conditions is to go to the cathode). It should be noted that cells were able to keep this new conditioned response for long periods of their cellular cycle, forgetting it later. Finally, we have quantitatively analysed the behaviour of these conditioned cells by studying the intensity of the locomotion responses, the directionality persistence, the persistence times, and several kinematic properties of the cell migration trajectories under conditioning such as the total distance traveled, the directionality ratio and the average speed. This exhaustive analysis unequivocally showed that the cellular conditioned behaviour is robust, and according to the parameters here analysed, these three species presented a similar motion structure characterised by the presence of conditioning for long periods, strong straightness in their trajectories, and high directional persistence.

The organisms analysed in our study are free-living naked lobose amoebae, the freshwater predators, belonging to the large and diverse supergroup *Amoebozoa*. Despite the seeming “simplicity” of these unicellular eukaryotes, the studies performed in *Amoeba* became a necessary basis for understanding various processes and phenomena, which are in the focus of attention of modern cellular and molecular biology, even if they are performed in organisms completely unrelated to protists phylogenetically distant to *Amoebozoa*. Thus, data on their locomotion mode, feeding behaviour, and cell cycle features contributed to the understanding of the structure and function of metazoan (including human) cells, also those of cancer^3–6^.

The amoebae’s life is a life in motion; they crawl over the substrates forming large or small protrusions searching for food and moving in response to external stimuli. As a single-celled organism that does not have complex structures responsible for the perception of external signals, *Amoebae* possess photo-, geo-, galvano- and chemotaxis^7–9^. Their locomotion’s type, known as “amoeboid movement”, is the most common locomotion mode among eukaryotic cells. Cells of multicellular organisms, including vertebrates, demonstrate a similar pattern of locomotion. Since the beginning of the 20^th^ century, several hypotheses have been proposed in an attempt to explain it (reviewed in^10^); nevertheless, the essential mechanisms of amoeboid movement remain to be elucidated.

Initial studies of the phagocytic ability of eukaryotic cells were preceded by investigations in *Amoeba*. Later, this activity was characterized in cells of the immune system of multicellular organisms, which are capable to engulf hostile cells. Interestingly, cancer cells also showed a clear feeding behaviour primarily oriented against neighboring cells, live or dead^11, 12^.

*Amoeba* is an obligatory agamic organism that demonstrates a very special cell cycle, during which a strategy of the so-called cyclic polyploidy (alternation of polyploidization and depolyploidization stages preceding reproduction) is implemented^13^. Such cycles of ploidy were described in cell cycles of some other unicellular eukaryotes. An analogous strategy is observed in mammalian cancer cells. Their similarity to unicellular organisms, specially amoebae, is considered a part of the atavistic theory of carcinogenesis, which suggests that cancer cells switch to a “selfish” lifestyle like unicellular organisms^6,14^.

In the present work, three species of large free-living amoebae were used as models. Two species belong to the genus *Amoeba, A. proteus* (Carolina Biological Supply Company, Burlington, NC. Item #-131306), and *A. borokensis* (Amoebae Cultures Collection of Institute of Cytology (ACCIC), St. Petersburg, Russia), and one third species belong to the genus *Metamoeba* – *M. leningradensis* (Culture Collection of Algae and Protozoa (CCAP), Oban, Scotland, UK, CCAP catalog number 1503/6).

The migratory trajectories of these three species have been here studied using an appropriate direct-current electric field (galvanotaxis)^8,15^ and a specific peptide (nFMLP, typically secreted by bacteria) as a chemoattractant (chemotaxis)^16^. More specifically, we have used a controlled electric field as the conditioned stimulus, and a specific chemotactic peptide as the unconditioned stimulus. Likewise, we have also carried out different control tests which indicated that cells independently exposed either to galvanotaxis or chemotaxis did not show any conditioned behaviour. Important qualitative and quantitative aspects of cellular migration under different guidance conditions have been addressed in this study. So, after the induction process (simultaneous galvanotaxis and chemotaxis), the three species exhibited the capacity to generate a new type of systemic motility pattern that prevailed for 40 minutes on average, which seems to be a strong evidence of a primitive type of associative memory in several species of unicellular organisms.

In parallel to our Pavlovian-like experiments, and to understand the dynamic characteristics of the locomotion movements in the migration of *Amoeba proteus*, we have recently analyzed also the locomotion trajectories of enucleated and non-enucleated amoebae using advanced non-linear physical-mathematical tools and computational methods^17^.The results showed that both cells and cytoplasts displayed migration trajectories characterized by non-trivial long-range correlations with periods of about 41.5 minutes on average, which corresponded to non-trivial dependencies of past movements. It is worth noting that this long-term memory (non-trivial correlations with 41.5 minutes on average), both in enucleated and nucleated cells, coincides with the Pavlovian-like dynamic memory (40 minutes on average) that we have detected now.

The Pavlovian-like experiments presented here were originally conceived by our research group after performing different physical-mathematical analyses of complex self-organized metabolic networks^18,19^. More specifically, using advanced tools of Statistic Mechanics and techniques of Artificial Intelligence, we could verify from computational approaches that Hopfield-like dynamics characterised by associative memory can emerge in these metabolic networks^19^. In this way our study showed quantitatively for the first time, that an associative memory is possible in unicellular organisms. This type of emergent property would be the manifestation of the complex dynamics underlying the functionality of the cellular metabolic networks at the systemic level. Such associative memory seems to correspond to an epigenetic type of cellular memory^20^.

In the present work, the results of our quantitative study were compared to other experimental findings in cellular behaviour with complex responses in bacteria, protozoa, fungus-like organisms and metazoans, which allow us to conclude that associative conditioning might be a widespread characteristic of unicellular organisms.

The cellular capacity of learning and therefore acquiring new adaptive systemic patterns of response to changes in the external medium could represent a fundamental process for cellular adaptation as an evolutionary mechanism through which unicellular organisms increase their fitness in their respective habitats.

In short, essential aspects of cellular systemic behaviour have been addressed in the present study. For a wide range of unicellular organisms, from prokaryotes to eukaryotes, cellular migration is an essential and tightly regulated systemic process which is involved in the major developmental stages of all complex organisms such as morphogenesis, embryogenesis, organogenesis, adult tissue remodeling, wound healing, immune cell activities, angiogenesis, tissue repair, cellular differentiation, tissue regeneration as well as in a myriad of pathological conditions^21^. However, how cells efficiently regulate their locomotion movements is still unclear. A novel knowledge of the control of cellular systemic behaviour in unicellular organisms such as cellular conditioning can define a new framework in the understanding of the mechanisms underlying the complex processes involved in the adaptive capacity of cells to the microenvironment. Moreover, such finding may constitute a significant advance in the comprehension of the biological responses involved in the evolutionary mechanisms for cellular adaptation, and in critical issues for human life like embryogenesis, tissue repair, and carcinogenesis^21^.

## Results

All the experiments were performed in a specific set-up (Fig. S1 and S2) consisting of two standard electrophoresis blocks, a power supply, two agar bridges and a structure made from a standard glass slide and covers (the experimental glass chamber) commonly used in Pathology-Cytology Laboratories (see details in the Methods section).

The first electrophoresis block was directly plugged into the power supply (with a direct electric field of about 300-600 mV/mm) while the other was connected to the first one via the two agar bridges, which allowed the current to pass through and prevented the direct contact of both anode and cathode with the medium where the cells will be placed later. Adequate control of intensity and voltage was established (Methods section and Fig. S3). In the center of the second electrophoresis block we placed the experimental chamber which consisted of a glass structure (described in the Methods section) and two glass sliding lateral pieces which could be displaced in the longitudinal direction. This way, when the sliding pieces were closed an inner laminar flux was produced in the chamber and, when they were open, the placement and collecting of the cells were possible, under the central piece made of a coverslip glass.

The experimental chamber not only allowed the electric current to pass through, but it also generated a nFMLP peptide gradient that the amoebae were able to detect and respond to. Therefore, the set-up is capable of exposing the amoebae to both stimuli simultaneously (galvanotaxis and chemotaxis) or independently of each other. The establishment of the nFMLP gradient was confirmed by measurement of the fluorescein-tagged peptide concentration with a plate reader (Fig. S5). The concentration of peptide in the middle part of the glass experimental chamber (where the amoebae were placed) increases quickly following the flow establishment (in 2 minutes the concentration rises from zero to approximately 0.2 mM), and this concentration further increases to 0.6 μM for at least 30 minutes.

Cells were placed in the middle of the experimental chamber and their displacements were monitored in small groups (see Methods section), being the individual trajectories recorded using a camera connected to a microscope. The digitized locomotion trajectories were analyzed in the form of time series. All the experiments were carried out in Chalkley’s medium, a standard nutrient-free saline medium, at ambient temperature.

### 1. Cellular displacement in the absence of stimuli

First, we observed that the migration of 169 individual cells belonging to the three species, *Amoeba proteus, Metamoeba leningradensis* and *Amoeba borokensis*, under the absence of stimuli, exhibited random locomotion patterns through which the amoebae explored practically all the directions of the experimentation chamber without a clear preference (Fig. 1a). To quantitatively analyse the directionality of the cells, we calculated the displacement cosine of each trajectory, where values close to −1 indicate a preference towards the left, and values close to 1 suggest preference towards the right. Since values did not follow a Gaussian distribution, values have been reported as median/interquartile-range, and non-parametric tests were applied to compare the data. In this experiment, the values ranged between −1 and 1 (*A. proteus*: 0.337/1.348, median/IQ; *M. leningradensis*: −0.431/1.194, median/IQ; *A. borokensis*: 0.173/1.422, median/IQ) confirming that in the absence of stimulus all cells moved randomly without any defined guidance. Experimental data: *A. proteus*, total cells 57, experimental replicates 8, number of cells per replicate 7-8; *M. leningradensis*, total cells 62, experimental replicates 8, number of cells per replicate 7-8; *A. borokensis*, total cells 50, experimental replicates 6, number of cells per replicate 7-10.

**Figure 1.**
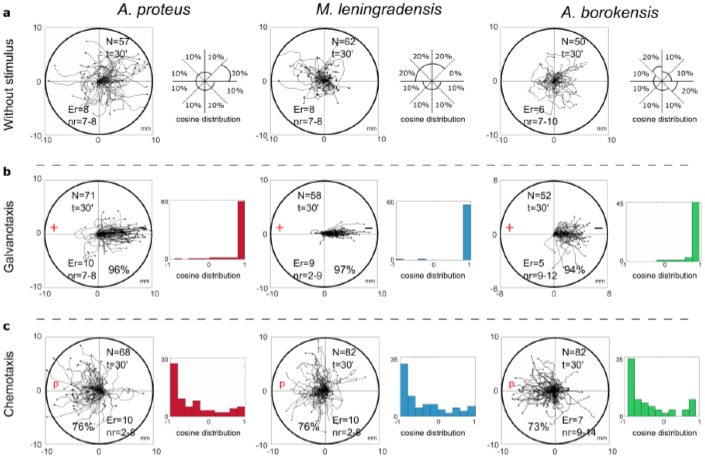
Migration trajectories of the three species under external experimental conditions. Panel **a** shows the migration without any stimuli by each of the three species (*A. proteus, M. leningradensis*, and *A. borokensis*, respectively). Practically, the amoebae explored all the directions of the experimentation chamber. In addition to the trajectories, the approximated percentage of cells that displaced in certain portion of the exploration chamber is represented. The space has been divided in 8 sections (thus π/4 angle amplitude was used for determining the regions). In panel **b**, the locomotion trajectories during galvanotaxis are depicted (96% of *A. proteus*, 97% of *M. leningradensis*, and 94% of *A. borokensis* migrated towards the cathode). Next to the trajectories, the respective histograms of the displacement cosines are illustrated. Finally, panel **c** represents the migration during chemotaxis. 76%, 76% and 73% of the respective cells migrated towards the chemotactic gradient. Alongside the trajectories, the histograms for the angles of each trajectory have been illustrated. “N” is the total number of cells, “Er” is the experimental replications, “nr” is the number of cells per replication, “t” time of galvanotaxis or chemotaxis, “p” chemotactic peptide (nFMLP), “+” anode, “-” cathode. Both the x and y axes show the distance in mm, and the initial location of each cell has been placed at the center of the diagram.

### 2. Cell locomotion in an electric field (galvanotaxis)

To study the cellular migration under galvanotaxis conditions, a controlled external direct-current electric field of about 300-600 mV/mm was applied to 181 amoebae belonging to the three species and analyzed in small groups (see “experimental data” below). The results showed that practically all the cells (96% of them) moved towards the cathode, indicating a strong directionality when the galvanotactic stimulus was applied (Fig. 1b). Specifically, 96% of *A. proteus*, 97% of *M. leningradensis* and 94% of *A. borokensis* migrated to the cathode. The values of the displacement cosines ranged between −1 and 1, (*A. proteus*: 0.984/0.068, median/IQ; *M. leningradensis*: 0.994/0.02, median/IQ; *A. borokensis*: 0.97/0.097, median/IQ) which confirmed that a fundamental behaviour characterised by an unequivocal directionality towards the cathode had emerged under these galvanotactic conditions (for more details, see Table 1). Next, the distributions of the values for the displacement cosines under galvanotaxis were compared to the values obtained in the experiment without stimulus, using Wilcoxon ran-sum test, which indicated that both behaviours were significantly different for the three species, and that this galvanotactic cellular behaviour is extremely unlikely to be obtained by chance (p-values: 10^−9^, 10^−18^ and 10^−10^; Z: −6.046, −8.78 and −6.27 for *A.proteus, M.leningradensis*, and *A.borokensis*, respectively). Experimental data: *A. proteus*, total cells 71, experimental replicates 10, number of cells per replicate 7-8; *M. leningradensis*, total cells 58, experimental replicates 9, number of cells per replicate 2-9; *A. borokensis*, total cells 52, experimental replicates 5, number of cells per replicate 9-12.

**Table 1.**

| Organism | Experiment | N | Displacement cosine (median/IQ) | Successfully responding (%) |
| --- | --- | --- | --- | --- |
| *A. proteus* | Galvanotaxis | 71 | 0.984/0.07 | 96% to (-) |
|  | Chemotaxis | 68 | -0.657/0.83 | 76% to P |
|  | Induction | 155 | 0.045/1.54 | 49% to P (+) |
|  | Conditioning | 76 | -0.84/1.02 | 75% to (+) |
| *M. leningradensis* | Galvanotaxis | 58 | 0.994/0.02 | 97% to (-) |
|  | Chemotaxis | 82 | -0.656/0.92 | 76% to P |
|  | Induction | 134 | -0.2/1.71 | 55% to P (+) |
|  | Conditioning | 65 | -0.908/0.45 | 82% to (+) |
| *A. borokensis* | Galvanotaxis | 52 | 0.97/0.10 | 94% to (-) |
|  | Chemotaxis | 82 | -0.632/0.90 | 73% to P |
|  | Induction | 169 | -0.332/1.69 | 60% to P (+) |
|  | Conditioning | 76 | -0.905/0.59 | 79% to (+) |

### 3. Cell migration under chemotactic gradient (chemotaxis)

Here, we analysed the trajectories of 232 cells belonging to the three species, which were exposed for 30 minutes to an nFMLP peptide gradient placed on the left side of the set-up (Fig. 1c). In this case, 75% of all cells migrated towards the attractant stimulus, the peptide (see Table1). Specifically, 76% of *A. proteus*, 76% of *M. leningradensis* and 73% of *A. borokensis* migrated to the peptide. The values of the displacement cosines were −0.657/0.827, median/IQ for *A. proteus*, −0.656/0.919, median/IQ for *M. leningradensis* and −0.632/0.9, median/IQ for *A. borokensis*. Since the medians for the three cosines of displacement were negatives, it was corroborated that most of the cells tended to migrate towards the left side, where the peptide was placed. The comparison between the displacement cosines under chemotaxis and galvanotaxis (p-values: 10^−18^, 10^−21^ and 10^−17^; Z: 8.74, 9.37 and 8.4 for *A. proteus, M. leningradensis*, and *A. borokensis*, respectively) corroborated that the systemic locomotion behaviour under the chemotactic gradient was totally different with respect to the observed under an electric field. Experimental data: *A. proteus*, total cells 68, experimental replicates 10, number of cells per replicate 2-8; *M. leningradensis*, total cells 82, experimental replicates 10, number of cells per replicate 2-8; *A. borokensis*, total cells 82, experimental replicates 7, number of cells per replicate 9-14.

### 4. Migratory behaviour under simultaneous galvanotactic and chemotactic stimuli (induction process)

After studying the behaviour of the three species without stimulus, under galvanotaxis and chemotaxis conditions, we exposed 458 cells in total to simultaneous galvanotactic and chemotactic stimuli for 30 minutes (induction process). The nFMLP peptide was placed on the left, in the anode (Fig. 2a-c). This experiment showed that 55% of all the amoebae moved towards the peptide-anode, while the remaining 45% migrated towards the cathode (Table 1). Specifically, 49% of *A. proteus*, 55% of *M. leningradensis* and 60% of *A. borokensis* migrated to the peptide-anode. The displacement cosines ranged from −1 to 1 (0.045/1.543, median/IQ) for *A. proteus*, (−0.2/1.706, median/IQ) for *M. leningradensis*, and (−0.332/1.685, median/IQ) for A. *borokensis*. This analysis quantitatively verified that two main cellular migratory behaviours had emerged in the experiment, one towards the anode and another towards the cathode (Fig. 2d-f). The statistical analysis (Wilcoxon rank-sum test) confirmed the presence of these two different behaviours for *A. proteus* (p-value=10^−27^; Z=10.74), *M. leningrandensis* (p-value=10^−23^; Z=9.93) and *A. borokensis* (p-value=10^−28^; Z=11.008). Experimental data: *A. proteus*, total cells 155, experimental replicates 21, number of cells per replicate 7-8; *M. leningradensis*, total cells 134, experimental replicates 18, number of cells per replicate 7-8; *A. borokensis*, total cells 169, experimental replicates 22, number of cells per replicate 5-11.

**Figure 2.**
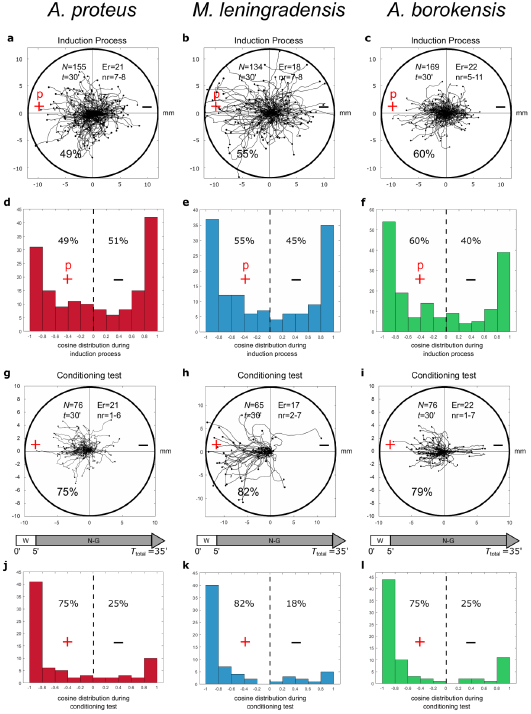
Induction processes and conditioning test. Panels **a**, **b**, and **c**, show the cellular migration under simultaneous galvanotactic and chemotactic stimuli (induction process) for *A. proteus, M. leningradensis*, and *A. borokensis*, respectively. Panels **d**, **e**, and **f**, illustrate the histograms of the displacement cosines from **a**, **b**, and **c** respectively. In panels **g**, **h**, and **i**, the trajectories under galvanotactic conditions of those cells that had previously migrated towards the anode-peptide is represented. It can be observed that the 75%, 82% and 79% of *A. proteus, M. leningradensis* and *A. borokensis* cells migrated towards the left positive pole, where the peptide was absent. Panels **j**, **k**, and **l**, illustrate the histograms of the displacement cosines from **g**, **h**, and **i** respectively. “N” is the total number of cells, “Er” is the experimental replications, “nr” is the number of cells per replication, “t” time of galvanotaxis or chemotaxis, “p” chemotactic peptide (nFMLP), “+” anode, “-” cathode. Both the x and y-axes show the distance in mm, and the initial location of each cell has been placed at the center of the diagram.

### 5. Emergence of cellular conditioned behaviour

To test if the cells that moved towards the anode during the induction process, presented some kind of associative conditioning in their migratory trajectories, we performed a conditioned behaviour test (Fig. 2g-i). For such a purpose, those cells that had previously migrated towards the anode-peptide during the exposition to the two simultaneous stimuli, were manually extracted and placed in a Petri dish with a normal culture medium (Chalkley’s medium) for 5 minutes, without any external stimuli, and then, they were re-exposed, for the second time, to the same single electric field, but without peptide. Note that when amoebae were placed in an electric field for long periods (30 minutes during the induction process), the probability of dying or, at least, detaching from the substrate and adopting a spherical shape increases sharply; therefore, after the induction process the cells were physically extracted and replated for 5 minutes to minimize cell damage. Next, we placed the cells in a new identical setup that had never been in contact with the peptide nFMLP, and filled it with clean Chalkley’s medium. In this context, the cells were again exposed to galvanotaxis for another 30 minutes.

Under these conditions, the analysis of the individual trajectories of 217 amoebae showed that most of the cells (78%) ran to the anode where the peptide was absent (Fig. 2g-i, and Fig. 3). Specifically, 75% of *A. proteus*, 82% of *M. leningradensis* and 79% of *A. borokensis* migrated to the positive pole. The fact that the majority of cells moved towards the anode in the absence of peptide corroborated that a new locomotion pattern had appeared in the cells (note that without the induction process, practically all the cells migrated towards the cathode under galvanotactic conditions).

**Figure 3.**
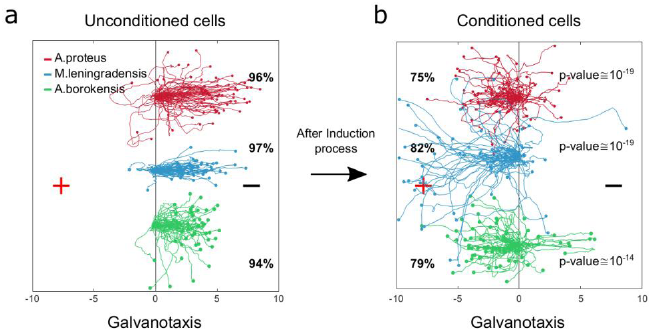
Comparison of amoebae migrations under two galvanotaxis conditions, before and after being conditioned. In red *Amoebae proteus*, in blue *Metamoeba leningradensis*, and in green *Amoebae borokensis*. **a** When galvanotactic stimulus was applied, practically all the cells of the three species migrated towards the cathode. **b** After induction process (simultaneous galvanotactic and chemotactic stimuli for 30 minutes, see Fig. 2) a new locomotion pattern had appeared in the cells, and the majority of amoebae moved towards the anode in the absence of peptide (75%, 82%, and 79% of the *A. proteus, M. leningradensis* and *A. borokensis* respectively). The p-values obtained when the displacement cosines of galvanotaxis and conditioning tests for each species were depicted, these values indicated that this newly acquired cellular behaviour is extremely unlikely to be obtained by chance. The three species responded similarly, suggesting the presence of associative conditions in most of them (78% on average). “+” indicates anode, while “-” cathode. The x-axis shows the distance in mm.

In this experiment the displacement cosines were ranged between −1 and 1, with −0.84/1.02 (median/IQ) for *A. proteus*, −0.908/0.447 (median/IQ) for *M. leningradensis*, and −0.905/0.588 (median/IQ) for *A. borokensis*, thus verifying that the majority of the amoebae displayed a new locomotion pattern characterized by movement to the anode during a galvanotactic stimulus without peptide (Fig. 2j-l). The results recorded in this experiment were compared to the ones obtained in the galvanotaxis without previous induction, and the test indicated that this newly acquired cellular behaviour is extremely unlikely to be obtained by chance (p-values: 10^−19^, 10^−19^ and 10^−14^; Z: 8.99, 8.91 and 7.52 for A. *proteus, M. leningradensis* and *A. borokensis*, respectively). Additionally, the comparison of all the cells from the conditioning tests against the galvanotactic responses was made, highlighting that this behaviour was extremely unlikely to be obtained by chance (p-value: 10^−49^, Z=14.75). Experimental data: *A. proteus*, total cells 76, experimental replicates 21, number of cells per replicate 1-6; *M. leningradensis*, total cells 65, experimental replicates 17, number of cells per replicate 2-7; *A. borokensis*, total cells 76, experimental replicates 22, number of cells per replicate 1-7.

### 6. Control tests

Different types of controls were established to complete the analysis of the previously mentioned experimental observations (Fig.4). First, to test the possible capacity of nFMLP peptide to change the migration of cells in an electric field, we have simultaneously exposed 55 *Amoeba proteus* to galvanotactic and chemotactic stimuli for 30 minutes, but in this case, placing the nFMLP peptide in the cathode (now in the left part, Fig. 4a). Afterwards, the same 55 amoebae were subjected to a single electric field, without peptide, for 30 minutes. Under the simultaneous exposition to the peptide and the electric field (placing the nFMLP peptide in the cathode), 93% of the cells migrated to the negative pole, while the remaining 7% exhibited a stochastic behaviour without a clear directionality towards any pole. Under the galvanotactic control, we observed that 98% of the cells ran towards the cathode. Therefore, the previous exposition of cells to the peptide nFMLP in an electric field (placing the nFMLP peptide in the cathode) did not induce any cellular behaviour characterized by movement towards the anode under galvanotactic conditions (total cells: 55, experimental replicates: 7, number of cells per replicate: 7-9).

**Figure 4.**
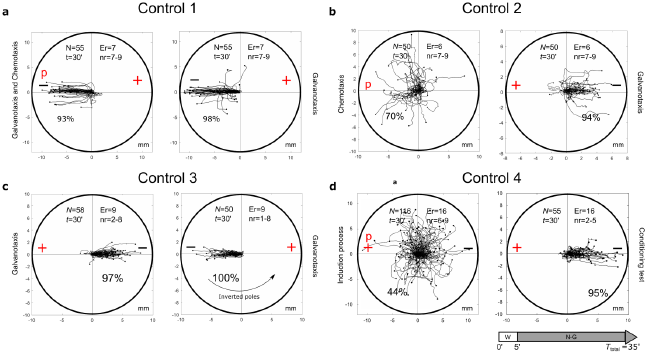
Control tests. **Control test on the capacity of nFMLP to change the migration of *Amoeba proteus* in an electric field**. A total of 55 *Amoeba proteus* were exposed simultaneously to galvanotactic and chemotactic stimuli for 30 minutes, placing the nFMLP peptide in the cathode. The same 55 amoebae previously exposed simultaneously to galvanotactic and chemotactic stimuli were subjected to a single electric field without peptide for 30 minutes. In both experiments, no cell showed clear directionality towards the anode. **Chemotactic and subsequent galvanotactic control**. A total of 50 *M. leningradensis* were exposed to nFMLP peptide for 30 minutes, and 72% of them migrated towards the anode. Then, those 50 cells were subjected to a galvanotactic stimulus during 30 minutes, and 90% of the cells migrated towards the cathode, therefore exhibiting normal galvanotaxis confirming that the exposure to the peptide did not alter the normal galvanotactic response. **Galvanotactic control with inverted polarity**. A total of 58 *M. leningradensis* were exposed to a galvanotactic stimulus for 30 minutes; next, the cells were exposed to another identical electric field with inverted polarity **b** (experimental replicates: 9, number of cells per replicate: 1-8). As it can be observed, practically all the cells showed a normal galvanotactic behaviour in both occasions. **Induction process and subsequent conditioning test of non-induced cells**. A total of 55 *M. leningradensis* were exposed simultaneously to the electric field and to nFMLP peptide for 30 minutes, and 44% of them migrated towards the anode. Then, those cells that migrated to cathode (56%) were subjected to a galvanotactic stimulus during 30 minutes, and 95% of such cells migrated towards the cathode, exhibiting normal galvanotaxis, and confirming that the previous exposure to the peptide did not alter the normal galvanotactic response. “N” is the total number of cells, “Er” is the experimental replicates, “nr” is the number of cells per replicate, “t” galvanotaxis time, “+” anode, “-” cathode, “P” nFMLP peptide. Both the x and y-axes show the distance in mm, and the initial location of each cell has been placed at the center of the diagram.

To test the cells that move towards the cathode during chemotaxis, 50 *M. leningradensis* were exposed to nFMLP peptide for 30 minutes and 72% of them migrated towards the peptide. Then, these 50 cells were subjected to a galvanotactic stimulus for 30 minutes (Fig. 4b). The results showed that 90% of the cells migrated towards the cathode, therefore exhibiting normal galvanotaxis. This experiment confirmed that the exposure to the peptide did not alter the normal galvanotactic response of cells (total cells: 50, experimental replicates: 6, number of cells per replicate: 7-9).

Next, a galvanotactic control of cells that moved towards the cathode was also performed. For such a purpose, 58 *M. leningradensis* were exposed to a galvanotactic stimulus for 30 minutes, next they were placed in a Petri dish filled with Chalkley’s medium for 5 minutes, and finally exposed to another identical electric field with inverted polarity (Fig. 4c). The results of this experiment showed that all the amoebae exhibited normal galvanotactic behaviour on both occasions. By inverting the position of the electric field, we demonstrated that the amoebae did not migrate to a specific point in the space (total cells: 58, experimental replicates: 9, number of cells per replicate: 1-8).

Finally, we tested the cells that moved towards the cathode during the induction process (simultaneous galvanotactic and chemotactic stimuli for 30 minutes) Fig. 4d. For such a purpose, 55 *M. leningradensis* that migrated towards the cathode during the induction process were again subjected to a controlled electric field (galvanotaxis for 30 mins). The results showed that practically all the amoebae (95%) migrated towards the cathode, confirming that these cells were unconditioned, and that their habitual behaviour, that is, to go to the cathode, was not altered by the induction process (total cells: 116, experimental replicates: 16, number of cells per replicate: 6-9).

### 7. Intensity of response in the conditioned cells

Here, we have analyzed the displacement module (measured in mm) of the conditioned cells which showed the intensity of the response in those cells that modified their migratory behaviour by the association of stimuli (Fig. 5a-b). Such a module corresponds to the vector, starting in the origin of the migratory trajectory and finishing at the end of it.

**Figure 5.**
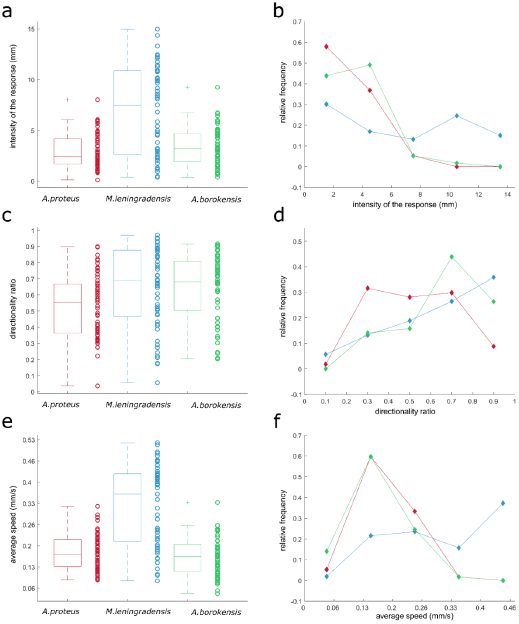
The intensity of response, directionality ratio, and average speed of the cells that acquired conditioned behaviour. **a**, **c** and **e** panels illustrate boxplots for the intensity of response (measured in mm), the directionality ratio, and the average speed (measured in mm/s) respectively (where *A. proteus* is represented in red, *M. leningradensis* in blue, and *A. borokensis* in green). On the side of each box, the values of the statistic for each cell were depicted by circles. Panels **b**, **d**, and **f** illustrate the frequency polygons corresponding to the statistics in panels **a**, **c,** and **e**, respectively. Here, the x-axis indicates the value of the statistic, while the y-axis represents the relative frequency (i.e., the number of cases divided by the total number of cells of that species) for the three cell types.

The results showed that the displacement module for *A. proteus*, values ranged from 0.103 to 8.017 (2.445/2.457, median/IQ), for *M. leningrandensis* was comprehended between 0.368 and 14.96 (7.5/ 8.26, median/IQ), and for *A. borokensis*, from 0.3701 to 9.229 (3.232/2.706, median/IQ). Kruskal-Wallis test indicated that there were significant differences regarding the intensity of response (p-value=10^−6^, χ^2^ statistic= 23.83), and therefore, pair-wise comparisons were made by using Wilcoxon rank-sum test. The obtained results indicated that significant differences were found between the values of *A. proteus* and *M. leningradensis* (p-value=10^−6^, Z=−4.468), as well as for the values of *A. borokensis* and *M. leningradensis* (p-value=0.00013, Z=3.81) but not between *A. borokensis* and *A. proteus* (p-value=0.2084, Z= −1.258). In Fig. 5a-b, the distribution of these statistics by boxplots and frequency polygons, respectively is depicted. Clearly, *M. leningradensis* responded much more intensively than the other two species, which exhibited a similar intensity in the conditioned responses. However, the values of metamoebae cells were more heterogeneous than those obtained from the other species, indicating more variability in its migratory conditioned behaviour; *A. proteus* presented the most stable intensity of the conditioned responses, with the lesser variability. Additionally, the response intensity pattern followed by *M. leningradensis* was very different from the observed in the other two species (Fig. 5b). The total number of conditioned cells analyzed was 167, and the experimental time 30 min.

### 8. Straightness of the conditioned cell trajectories

To analyse the straightness in the direction followed by the conditioned cell we have calculated the directionality ratio (Fig. 5 c-d). This statistic test quantifies the trajectory straightness, which ranged between 0 (for fully curved trajectories) and 1 (for fully straight tracks). First, we computed the total trajectory length, next, we obtained the euclidean distance between the start point and the endpoint, and finally, we defined the directionality ratio as the quotient between these two figures. The results showed that for *A. proteus*, values ranged between 0.0362 and 0.899 (0.555/ 0.304 median/IQ), for *M. leningradensis* from 0.056 to 0.970(0.688/0.412, median/IQ), and for *A. borokensis*, from 0.2032 to 0.916 (0.681/ 0.304 median/IQ). In this case, Kruskal-Wallis test indicated that at least the directionality ratio of one of the species was significantly different (p-value=0.002, χ^2^ statistic= 12.66), and therefore, pair-wise comparisons were made. The obtained results showed that no significant differences were found between the directionality ratio values of *A. borokensis* and *M. leningradensis* (p-value=0.697, Z=0.3888), however the values of *A. proteus* were significantly different to the values of *M. leningradensis* (p-value=0.0055, Z=−2.775), and *A. borokensis* (p-value=0.0008, Z= −3.34). This data suggested that *A. borokensis* and *M. leningradensis* followed similar and strong straightness in their conditioned migratory direction, going to the migratory target much more directly than *A. proteus* cells, which exhibited a more wandering trajectory in their movements. Interestingly, again, *M. leningradensis* showed more heterogeneity in their directionality values than the obtained from the other species, indicating more variability in their conditioned responses. Moreover, *A. borokensis* presented the largest number of cells with high straightness in their migratory movements (Fig. 5d). The total number of conditioned cells analyzed was 170, and the experimental time 30 min.

### 9. Average speed of the conditioned cells

Another kinematic property analysed was the average speed (measured in mm/s) of the conditioned cells. The results showed that the values for *A. proteus* ranged from 0.093 to 0.324 (0.173/ 0.084, median/IQ), for *M. leningrandensis* between 0.09 and 0.523 (0.362/ 0.212, median/IQ), and for *A. borokensis*, from 0.05 to 0.336 (0.167/0.086, median/IQ), see Fig. 5e-f. The test comparing the average speed of the three species indicated that the distributions were significantly different (p-value= 10^−11^, χ^2^ statistic= 48.43). The obtained results indicated that no significant differences were found between the speed values of *A. proteus* and *A. borokensis* (p-value=0.2918, Z= 1.054), but the average speed was significantly different between *A. proteus* and *M. leningradensis* (p-value=10^−9^, Z=−5.814), and between *A. borokensis* and *M. leningradensis* (p-value=10^−10^, Z=6.197). Conditioned *M. leningradensis* cells moved significantly faster than the other two species. However, these metamoebae cells migrated more dispersedly that *A. proteus* and *A. borokensis* cells, which in turn, showed a similar behaviour for the average speed (Fig. 5e). In the case of *A. proteus* and *A. borokensis* cells, the majority of speed values were concentrated around 0.16 mm/s (Fig. 5f). In addition, the distance traveled (measured in mm) was previously estimated to obtain the average speed. These results showed that the values for *A. proteus* ranged from 2.777 to 9.714 (5.18/ 2.523, median/IQ), for *M. leningradensis* varied between 2.692 and 15.7 (10.871/ 6.373, median/IQ), and for *A. borokensis*, from 1.485 to 10.073 (5.0022/2.5647, median/IQ). The total number of conditioned cells analyzed was 170, and the experimental time 30 min.

### 10. Directionality persistence level in the conditioned cells

Next, we studied the persistence of the movement by analyzing Spearman’s correlation coefficient relating the x coordinate of the displacement of the cells, and the respective time step (Fig. 6). Values close to −1 indicated that cells migrated persistently towards the left pole (anode), while values close to 1 corresponded to a strong tendency towards the right pole (cathode), and intermediate values suggested little persistence of migration to any pole. Results of our analysis indicated that values of *A. proteus* ranged between −1 and 0.171 (−0.934/ 0.184 median/IQ), of *M. leningrandensis* from −1 to 0.017 (−0.976/0.137, median/IQ), and of *A. borokensis*, from −1 to 0.663 (−0.99/ 0.058 median/IQ).

**Figure 6.**
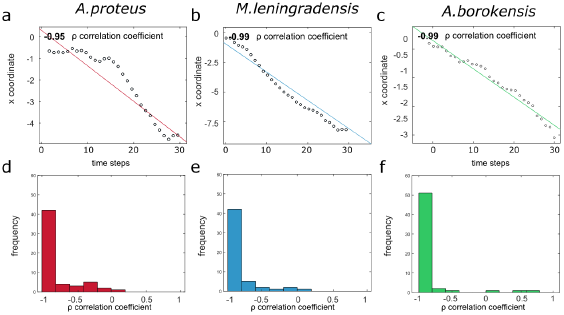
Persistence towards the anode for the cells that acquired conditioned behaviour. **a**, **b** and **c** panels illustrate Spearman’s correlation coefficient ρ relating the x coordinate and the time steps for three prototypical cells, each one of them corresponding to one of the species (*A. proteus, M. leningradensis* and *A. borokensis*). As it can be observed, there is a strong negative correlation, denoting that as time increases, cells tend to maintain their migratory direction towards the anode, which indicates high persistence during the displacement. **d**, **e,** and **f** panels represent histograms for Spearman’s correlation coefficients for the three species, respectively. It can be noticed that the majority of the cells present a strong negative correlation, corresponding to strong persistence towards the anode.

In this case, the test showed the presence of significant differences between the statistic values (p-value=0.0008, χ^2^ statistic= 14.18). The results indicated that no significant differences were found between the correlation values of *A. borokensis* and *M. leningradensis* (p-value=0.459, Z=0.739), however, the values of *A. proteus* were statistically different compared to those of *A. borokensis* (p-value=0.00016, Z= 3.769) and *M. leningradensis* (p-value=0.0127, Z=2.492). The results showed that the persistence values of *A. borokensis* were higher than those of *A. proteus*, but no differences were found when these values were compared to those obtained from *M. leningradensis*. All the amoebae analyzed exhibited a very strong preference towards the anode. To illustrate this behaviour, in Figure 6a-c we have represented three prototypical series and their correlation coefficient calculation. Also, in Figure 6d-f we represent through histograms all the correlation values obtained for the trajectories of the three species. The total number of conditioned cells analyzed was 170, and the experimental time 30 min.

### 11. Persistence time in the conditioned cells

The motility patterns acquired by amoebae are also characterised by a limited period of conditioned behaviour. Figure 7 c, d and e display how the trajectories of 15 amoebae that previously had acquired the systemic conditioned behaviour after the induction process gradually lost the persistence towards the anode and turned back to the cathode, under galvanotaxis, thus losing their acquired behaviour.

**Figure 7.**
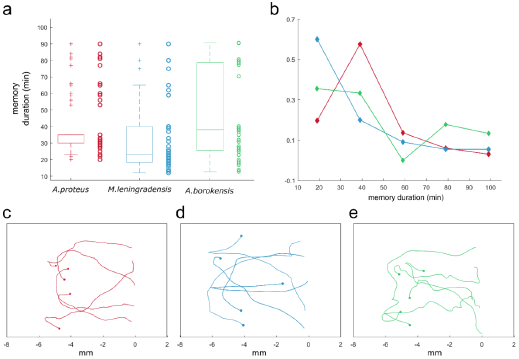
Persistence time for the cells that acquired conditioned behaviour. **a** Boxplot representing the duration of the acquired associative behaviour (up to 90 minutes) for the three species (*A. proteus* in red, *M. leningradensis* in blue and *A. borokensis* in green). On the side of each box, we depict the persistence time values with circles. **b** Frequency polygons representing the duration for each species. The x-axis indicates the duration in minutes, while the y-axis represents the relative frequency for the three cell types. As it can be observed, the majority of the cells present persistence times ranging between 20-50 minutes. Panels **c**, **d,** and **e** show how the trajectories of 15 prototypical amoebae (belonging to *A. proteus, M. leningradensis* and *A. borokensis*) after the induction process lost gradually the persistence towards the anode and turned back to the cathode, under new galvanotaxis conditions. Here, the x-axis indicates the distance traveled in mm.

We estimated the temporal persistence of the conditioned behaviour calculating the time (measured in minutes) during which these cells showed a directional response to anode during galvanotaxis after an induction process. For such a purpose, 166 amoebae (66 *A. proteus*, 55 *M. lenigradensis*, 45 *A. borokensis*) that had previously migrated towards the anode-peptide during the exposition to the two simultaneous stimuli (induction process) were manually extracted and placed for 5 minutes on a normal culture medium (Chalkley’s medium) in a small Petri dish in absence of stimuli. Next, the cells were deposited on a new identical set-up that had never been in contact with the chemotactic peptide nFMLP and exposed for the second time to a single electric field, without peptide, for 30 minutes. This process was repeated two more consecutive times. The time elapsed until the cells forgot the conditioned response, turned around, and returned to the cathode was estimated (Fig. 7) and the results indicated that the temporal persistence for *A. proteus*, ranged between 20 to 90 (35/5, median/IQ), for *M. leningradensis* between 12 and 90 (23/21.75, median/IQ), and from 13 to 90 (38/52.5, median/IQ) for *A. borokensis*.

The test indicated that at least the persistence times of one of the species was significantly different (p-value=0.0017, χ^2^ statistic= 12.7). Temporal values of *M. leningradensis* were found to be significantly different when compared to the values of *A. proteus* (p-value=0.0017, Z=3.134) and *A. borokensis* (p-value=0.0032, Z=−2.945), but no differences were found between the values of *A. proteus* and *A. borokensis* (p-value=0.466, Z= −0.728). The results suggested that the temporal persistence of conditioning in *M. leningradensis* cells is the smallest one and that the temporal values of *A. proteus* and *A. borokensis* were similar. However, the values of *A. borokensis* were much more dispersed and less homogeneous than those of *A. proteus* or *M. leningradensis*, which could imply that cells belonging to this species may have a wider range of persistence than the others. Fig. 7b shows that the patterns of loss of persistence towards the anode (loss of conditioning) were very similar in the three species. The whole analysis indicated that the total average time of the cells belonging to the three species that lost the acquired motility pattern was 39.58±22.48 (mean±sd) minutes. So, cells belonging to three species of unicellular organisms were able to acquire a new systemic behaviour from the association of stimuli; they were also able to keep it for long periods compared to their cellular cycle, and they forgot them later.

Finally, a violin graph (Fig. 8) depicts the most relevant results of our quantitative analysis. This figure shows the distributions of the response intensity (Fig. 8a), the directionality ratio (Fig. 8b), average speed (Fig. 8c), Spearman’s correlation coefficient (Fig. 8d), and temporal persistence (Fig. 8e) for conditioned amoebae belonging to *A. proteus, M. leningradensis* and *A. borokensis,* respectively. From this exhaustive analysis, we concluded that *A. proteus* cells presented less variability and more stability on their responses, characterised by strong persistence, less speed, and less distance traveled than the other two species. In addition, *A. proteus* cells were the ones that exhibited greater wandering in their migratory movements. Conversely, *M. leningradensis* were the less stable cells, presenting less homogeneity and more variability than the other two species. Additionally, *M. leningradensis* migrated faster than the other two species, but on the other hand, the temporal persistence of the conditioned behaviour in these cells was significantly smaller than the others.

**Figure 8.**
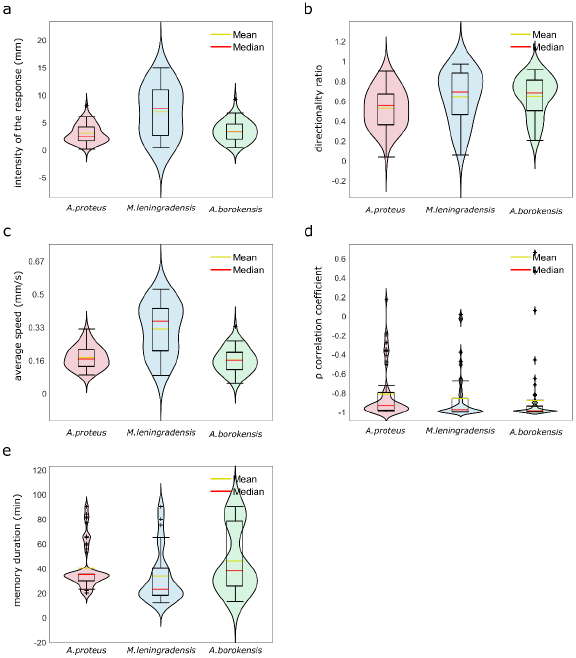
Violin plots for cells of the three species that acquired conditioned behaviour. In this graph, we represent by Violin plots the probability distribution, the boxplots, and the mean value of the five main quantitative characteristics. Panel **a** depicts the intensity of response (in mm), panel **b** illustrates the directionality ratio, panel **c** represents average speed (in mm/s), panel **d** depicts Spearman’s correlation coefficient between time and the x-axis of the trajectory, and **e** illustrates the persistence time of the acquired new behaviour (mins). *A. proteus* cells presented conditioned responses characterised by strong persistence, less speed and less distance traveled than the other species. *M. leningradensis* were the less stable cells i.e., demonstrated less homogeneity and more variability than the other species; additionally, this species showed more intensive and fast migration than the other two. *A. borokensis* cells exhibited more directionality persistence than the other two species, with high straightness towards the anode. The three species analysed show extremal values in their correlation coefficient (close to −1), and it can conclude that the cellular conditioning for the three species is a very robust behaviour with strong directional persistence.

*A. borokensis* were the cells which presented more directionality persistence than the other two species, with high straightness towards the anode. Regarding speed and intensity of response, *A. borokensis* behaved similarly to the *A. proteus* cells. Besides, the temporal persistence was the most disperse of the three species, and in average, the largest. We can conclude that the cellular conditioning was robust, and these three species presented a similar motion pattern structure, characterised by persistence times larger than 20 minutes, directionality ratios close to 0.6, and persistence directionalities close to −1, indicating the presence of conditioning for long periods, remarkable straightness in their trajectories and strong directionality persistence towards the anode pole, respectively.

## Discussion

Over centuries, the combined work of naturalists, philosophers, physiologists and scientists has shaped a very plentiful and venerable history of the essential principles that define the basic forms of associative memory and learning processes^22^. Among all of them, it is worth noting the Nobel Prize Laureate Ivan Pavlov who was the first to establish the fundaments of the associative memory in his classic studies with dogs^23^. Following a similar conceptual framework of Pavlov’s experiments, and using an appropriate direct-current electric field (galvanotaxis) and a specific peptide (nFMLP, typically secreted by bacteria) as a chemoattractant (chemotaxis), we have addressed here essential aspects of the cellular systemic behaviour. The migration trajectories of more than 2000 individual cells belonging to three different species of unicellular organisms, *A. proteus, M. leningradensis* and *A. borokensis*, have been exhaustively studied in this work, confirming that these three species were able to modify their systemic response to a specific stimulus by associative conditioning.

First, we have observed that cell locomotion in absence of stimuli exhibited a random directional distribution by which amoebae explored practically all the directions of the experimentation chamber. Second, cells showed an unequivocal systemic response consisting of the migration to the cathode when exposed to a direct electric field (galvanotaxis). Third, amoebae were studied during chemical guidance by exposing them to an nFMLP peptide gradient. In this experiment, 75% of cells on average migrated towards the chemotactic stimulus showing stochastic locomotion movements with robust directionality towards the peptide (chemotaxis). All these unicellular organisms belonging to the three species here studied responded in concordance with other similar experimental observations, carried out in absence of stimuli^17^, in galvanotactic conditions^15,16^, and under a chemotactic stimulus^24^.

An essential question of contemporary biology is to know if cells are capable of acquiring new behavioural systemic patterns to adapt themselves to changes in the external medium. For such a purpose, those cells that had previously migrated towards the anode-peptide during the dual simultaneous exposition to galvanotactic and chemotactic stimuli (induction process) were exposed to a single electric field, without peptide. Under these conditions, most of the cells (78% on average) ran to the anode where the peptide was absent. The fact that the majority of induced cells moved towards the anode in the absence of peptide corroborated that a new locomotion pattern had appeared in the cells. Note that without the induction process practically all the cells migrated towards the cathode under galvanotactic conditions and, conversely, after induction process most of the cells went to the anode.

When the exposition to a stimulus related to the amoebae nourishment (the peptide nFMLP) is accompanied by an electric field, and the peptide is placed in the anode (induction process), some amoebae seemed to associate the anode with the food (the peptide). After the induction process, testing under galvanotactic conditions, most of the conditioned amoebae ran towards the anode, where the peptide was absent, in contrast to their known tendency to move to the cathode, which demonstrates their modified systemic conduct.

To better understand this cellular conditioning, we analysed a great number of individual conditioned amoebae by studying the intensity of the migratory movements, the directionality persistence, the total distance traveled, the average speed, the directionality ratio, and the persistence times.

This analysis has shown that *A. proteus* cells presented the most stable conditioned responses, with less variability, as well as exhibiting strong persistence, less distance traveled, and less speed of locomotion movements. The temporal persistence was high in these amoebae. However, they exhibited greater wandering in their migratory trajectories.

The conditioned *M. leningradensis* showed the maximum intensity of response during its locomotion movements but presented much less stable behaviour than *A. proteus* i.e., they exhibited greater variability and heterogeneity in all the tests performed. Also, these amoebae cells were those that traveled the most distance, exhibiting maximum speed in their movements, and sharp persistence with strong straightness. Their persistence times were the shortest of the three species.

Conditioned *A. borokensis* were the cells that showed greater persistence with high preference towards the anode as well as robust straightness. These amoebae exhibited similar distances traveled compared to the *A. proteus* with very close migration speeds and persistence times, but with greater dispersion and variability. Besides, their persistence times were the longest of the three species.

Cells belonging to the three species analyzed showed extremal values in their Spearman’s correlation coefficient (close to −1) i.e., there is very high directionality persistence during the migratory displacement of all amoebae. In fact, this is a key characteristic of conditioned cells. The strong negative correlation denoted that as time increases, cells sharply tended to maintain their migratory direction towards the anode during a large period (despite their usual behaviour is to move towards the cathode). Additionally, this magnitude is scale-invariant for the two variables considered, in this case, the x coordinate indicating the position and the time step t. This analysis showed that cellular conditioning was robust for the three species with strong directional persistence, and such a result represents a fundamental aspect of the behaviour of conditioned cells.

The kinematic variables studied (intensity of the migratory movements, the total distance traveled, the average speed, and the directionality ratio) highlighted the different responses that cellular life exhibited at individual level, even within the same species. However, we can conclude that the cellular conditioning for the three species was robust because these three species presented a similar motion migratory structure, characterized by persistence times larger than 20 minutes, directionality ratios close to 0.6, and persistence directionalities close to −1, indicating the presence of conditioning for long periods, remarkable straightness in their trajectories and strong directionality persistence towards the anode. Most conditioned cells moved towards the anode in the absence of peptide thus corroborating that a new locomotion pattern had appeared in the cells (specifically, 75% of *A. proteus*, 82% of *M. leningradensis*, and 79% of *A. borokensis* migrated to the anode). These cells were able to keep this new conditioned response for long periods of time with respect to their cellular cycle (39.58 ± 22.48 minutes), forgetting it later. The results were compared to those obtained in the galvanotaxis without previous induction, and all tests indicated that this newly acquired cellular behaviour is extremely unlikely to be obtained by chance (p-values: 10^−19^, 10^−19^ and 10^−14^ for *A. proteus, M. leningradensis*, and *A. borokensis*, respectively). As a matter of fact, the comparison of all the cells from the conditioning tests against the galvanotactic responses emphasized that it was extremely unlikely to obtain such conditioned behaviour by chance (*A. proteus, M. leningradensis, A. borokensis* p-value: 10^−49^). We can conclude that cellular conditioning represents a robust behaviour with very strong directional persistence in the three species.

On the other hand, all controls indicated that cells that were exposed independently to galvanotaxis or chemotaxis or under simultaneous galvanotactic and chemotactic stimuli (but placing the nFMLP peptide in the cathode) did not present any observable conditioned pattern. Pavlov described four fundamental types of persistent behaviour provoked by two stimuli. The experiments on cellular conditioning exposed here were based in one of them, the so-called “simultaneous conditioning”, in which both stimuli are applied at the same time. Nevertheless, strictly speaking we cannot classify our findings as the classical Pavlovian conditioning since not all controls for classical conditioning studies were performed yet^25^.

In parallel to our Pavlovian-like experiments, for better understanding the dynamic characteristics of the locomotion movements and to quantitatively study the role of the nucleus in the migration of *Amoeba proteus*, we have previously analysed the movement trajectories of enucleated and non-enucleated amoebae using advanced non-linear physical-mathematical tools and computational methods^17^. In this study, we analysed the relative move-step fluctuation along their migratory trajectories by applying the root mean square fluctuation, a classical method in Statistical Mechanics based on Gibbs and Einstein studies^26,27^ that has been developed and widely applied to quantify different time-series. The results showed that both cells and cytoplasts displayed migration trajectories characterized by non-trivial long-range positive correlations with periods of about 41.5 minutes on average in all the analysed cells and cytoplasts, which corresponded to non-trivial dependencies of the past movements. Therefore, each cellular move-step at a given point is strongly influenced by its previous trajectory. This dynamic memory (non-trivial correlations) represents a key characteristic of the movements of *Amoeba proteus* during cell migration^17^. It is worth noting that this temporal persistence (non-trivial correlations with 41.5 minutes on average) in both enucleated and nucleated cells matches with the Pavlovian-like dynamic memory (40 minutes on average) obtained here.

Our results unequivocally evidence that cellular conditioning is a robust and stable cellular property, and that the acquired systemic responses are very similar in the three species analyzed. The fact that unicellular organisms belonging to three different species such as *A. proteus, M. leningradensis*, and *A. borokensis* are able to associate external signals and consequently generate new robust migratory responses, which can be remembered for long periods, show that cellular conditioned behaviour could correspond to a widespread phenomenon in the cellular life.

The inkling of cellular behaviour consistent with associative conditioning has been observed in different kingdoms including bacteria, protozoa, fungus-like organisms and metazoan. For instance, gene expression patterns in *E. coli* are almost identical when responding to either a drop in oxygen concentration levels or to a rise in the ambient temperature. This behaviour is thought to be a consequence of the association of higher temperatures with an anaerobic environment, such as it is the prevailing context in mammalian intestinal tracts^28^. H. Armus and colleagues have identified in *Paramecium caudatum* that this protist is able to develop a preference for illumination level using a mild electric shock as a reinforce^29^. Fungus-like organisms such as *Phisarum polycephalum* has been shown to exhibit a primitive type of learning^30^. Moreover, this unicellular multinucleate plasmodium was observed to acquire new thermotactic behaviours when exposed to a temperature gradient and a source of nutrients. While under normal circumstances, *Phisarum polycephalum* tend to migrate toward higher temperatures up to 30°C, after conditioning, it shows a preference for cooler temperatures^31^. Recently, human pancreatic β cells have also been demonstrated to exhibit associative conditioning behaviours. In short, temporal potentiation of sensitivity to glucose was achieved by replacing glucose concentrations with other secretagogues in combination with carbachol, giving rise to a newly formed behaviour that could be explained by a short-term associative conditioning process^32^.

It is still too early to hypothesize about the molecular mechanisms supporting the cellular associative conditioning. In this sense, we want to point out that the Pavlovian-like experiments presented here were originally conceived in 2013 when we studied self-organized enzymatic activities arranged in dynamic metabolic networks^18^. From these studies, we could verify using advanced tools of Statistic Mechanics and techniques of Artificial Intelligence the emergence of Hopfield-like dynamics in self-organized metabolic networks which were characterized by exhibiting associative memory, similar to neural networks^19^. In that study, the associative memory emerges as a consequence of the complex metabolic dynamics that take place within the cell at systemic level. Consequently, our work quantitatively showed, for the first time, that an associative memory was also possible in unicellular organisms, and such type of memory would correspond to an epigenetic type of cellular memory^20^.

Behaving efficiently is an essential issue for adaptation. Throughout evolution, the organisms that adapt successfully their behaviour to the environment do increase their possibilities to survive and reproduce compared to those that cannot. Associative learning in animals has long been considered essential for the efficient adaptation and processing information about the environment ^33^. In free-living cells, unlike habituation and sensitization, associative conditioning results in responses that are reliably correlated with the state of the external medium, allowing unicellular organisms to recognize significant events and respond quickly and appropriately to specific signals. The association of these cues will benefit tracking current conditions and map them for optimal or near-optimal responses. This type of learning can lead to very rapid, rather than gradual, behavioural changes, that enable fast detection of correlated features of complex environments. All these elements, allow to consider cellular associative conditioning as a powerful optimizing mechanism to perform efficiently different behavioural sequences under variable ecological circumstances. Cellular associative conditioning would be a good example of how evolution can find elegant solutions to an adequate adaptation to the microenvironments.

In short, essential aspects of systemic cellular behaviour have been addressed here. In particular, we have exhaustively analysed several important characteristics of cellular migration, a systemic behaviour essential in cellular development, and the functional maintenance of both free-living cells and cells of multicellular organisms. In humans, embryogenesis, organogenesis, immune responses, and tissue repair, for instance, require very precise and complex migratory movements of cells. Any error in the control of these migratory processes may result in important consequences such as mental retardation, cardiovascular diseases, and cancer^34^. The metastatic process allowing tumor cells to abandon the primary tumor and migrate to distant organs is a leading cause of death in patients with cancer. A better knowledge of the processes that control such migration will contribute to reducing the cancer-associated mortality^21, 35^. This connection with medical practice increases the importance to expand our knowledge about cellular conditioning. Moreover, the cellular capacity to acquire new adaptive systemic patterns to face changes in the external medium could represent an evolutionary mechanism whereby cells become more capable of living in diverse habitats. Cells are continuously receiving different biochemical and bioelectric signals, and our experiments have evidenced that unicellular organisms are capable to associate these signals and generate new adaptive migratory behaviours that can be remembered for long periods. This finding might be essential to understand some main principles of cellular evolution and its adaptive strategies involved in enhancing their evolutionary fitness to external medium. Consequently, the observation of this new cellular property can help in understanding of many cellular processes difficult to explain in the current conceptual framework and can open a new perspective of research in the Sciences of Life.

## Methods

### Cell cultures

*Amoeba proteus* (Carolina Biological Supply Company, Burlington, NC.Item # 131306) were cultured alongside *Chilomonas* as food organisms at around 21 °C on Chalkley’s simplified medium (NaCl, 1.4 mM; KCl, 0.026 mM; CaCl2, 0.01 mM), (Carolina Biological Supply Company Item #131734) and previously baked wheat corns. *Metamoeba leningradensis* (Culture Collection of Algae and Protozoa, Oban, Scotland, UK, CCAP catalog number 1503/6) were cultured in the same conditions as *Amoeba proteus. A. borokensis* (Amoebae Cultures Collection of Institute of Cytology –ACCIC, St. Petersburg, Russia) were cultured in Prescott & Carrier’s media, and supplied by 0.5 ml of Chilomonas and Colpydium (ACCIC, St. Petersburg, Russia^5^) as food organisms twice a week; these two organisms were also cultured in Prescott & Carrier’s media^37^ with flamed rice grains.

### Experimental set-up

The experiments in this study were performed in a device shown in Figure S1 and Fig S2 that was composed of two blocks of electrophoresis, 17.5 cm long each (Biorad Mini-Sub cell GT), a power supply (Biorad powerbank s2000), two bridges of agar (2% agar in 0.5 N KCl, 10-12cm long), and a chamber made of glass covers and slides.

The first block was plugged to the power supply and connected to the second one by the two agar bridges this way preventing the direct contact of the second electrophoretic block with the power supply. The blocks were composed of three parts: two wells filled with the conductive medium (Chalkley’s simplified medium) and an elevated platform. The experimental glass structure was placed on the top of the elevated platform of the second block. This way, the implementation of a laminar flux was possible when the chamber was closed. Opening the chamber allowed to get it and extract the cells.

### Experimental chamber

Standard glass slides and covers (total 4 pieces) composed the experimental chamber, that is, a 75×25 mm slide and three small pieces resulting from a careful cut of three standard long 40×24×0.1 mm cover glasses (Fig. S1–S2).

#### 1. Preparation of the sliding components

Three one-use-only cover glasses were trimmed to get one central piece of 3×24×0.1 mm and two lateral sliding pieces of 40×24×0.1 mm each.

#### 2. Modified glass slide

This glass supports the central and sliding parts of the structure (experimental chamber) (Fig. S1–S2). We used silicone to glue two covers to a glass slide. This structure was left to dry for 24 hours. Then, the protruding portions of the two cover slides (60×20×0.1 mm) were trimmed, leaving two 60×4×0.1 mm glass strips, which act as the lateral walls of the experimental chamber.

#### 3. Mounting the experimental chamber on the set-up

The resulting modified glass slide was placed on the top of the elevated central platform of the second electrophoresis block. Under it, an oil drop was placed to prevent that the Chalkley’s simplified medium can pass under the glass structure. The oil must fill the entire surface of the glass structure so that no liquid can pass through. On the top of the modified slide the central piece and the two sliding lateral glasses were placed.

#### 4. Cell placement in the chamber

Amoebae were placed under the central piece of the experimental glass chamber in 30 μl of clean Chalkley’s simplified medium. The process must be performed rapidly; otherwise the amoebae will adhere to the tip of the pipette and damage their cell membrane.

#### 5. Laminar flux implementation

Cells were left to rest for around 2 minutes on the experimental chamber prior to every experiment. This way, they were allowed to adapt to the new environment, firmly attach to the glass surface and begin moving around the experimental chamber. The wells of both electrophoretic blocks were carefully filled with Chalkley’s simplified medium until reaching the base of the modified slide but not the sliding glasses. Then, the sliding glasses were pushed down with a pipette to put them in contact with the Chalkley’s simplified medium allowing the medium to sprawl by surface tension. Next, the two sliding glasses were gently moved alongside the longitudinal dimension of the experimental glass structure until touching the central glass piece to create a laminar flux.

#### 6. Addition of the peptide to the set-up

When required, always before the establishment of the electric field, 750 μl of 2×10^−4^ M nFMLP peptide solution was added to the Chalkley simplified medium (75 ml) in the well corresponding to the positive pole of the second electrophoretic block. The same volume (750 μl) of Chalkley’s medium was removed from this well before the peptide was added. Keeping the same volume of medium in both wells is necessary to avoid harmful medium flows through the experimental chamber, which would affect the amoebae’s behaviour.

#### 7. Cell extraction

Cells were rescued from the chamber by sliding the top lateral pieces of the glass structure using the tip of a micropipette.

### Cell preparation

*A proteus, M. leningradensis* and *A. borokensis* may show different physiological variations due to slightly differing culture conditions. Prior any experiments, cells to be studied were starved for 24 hours in clean axenic Chalkley simplified medium in the absence of any external stimuli. Only cells that were actively moving, did not display many thin or upwards pseudopodia and showed a spindled or oval shape were selected for the experiments.

All cells analyzed were washed in Chalkley simplified medium prior to experimentation and then placed under the central piece of the glass chamber. The glass chamber was never closed before all amoebae appeared to be firmly attached to substrate, what happened in about a minute on average. The experiments were always performed using no more than 10 cells. Consecutive experiments tended to use always a smaller number of cells, since only those cells showing a potentially conditioned behavior were tested further. Interestingly, *M. leningradensis* displayed more variability of behaviors and shapes. They were also more difficult to handle, showing less adherence to the glass and more tendency to stick to the pipette tip.

All experiments implying the use of galvanotaxis were traumatic for cells, and could alter their response to following experimentation. To minimize harmful effects, the amoebae must always be handled with extreme caution, only healthy and fit amoebae (as previously described in this section) shall be used and the electricity parameters’ value kept within the provided optimal ranges during experimentation.

### Electric field (galvanotaxis)

The electric field applied to the first electrophoresis block was conducted to the second by the two agar bridges. Direct measurements in the second block close to the experimental chamber made it possible to control and maintain the current between 58.5 and 60V (334-342 mV/mm).

To ensure that the intensity required to successfully perform the experiments was supplied, it is important that during construction of the experimental chamber that the longitudinal strips were ≤4 mm in width and ≥0.1 mm longer. Height of the longitudinal strips was adapted by modifying the amount of silicone used to glue the cover slides pieces to the glass slide while constructing the chamber. Additionally, in order to have an immediate control of the intensity of the current, a variable resistance of 1 megohm was installed in series, and after it a milliamperimeter (Fig. S3). During the experiments, by modifying the variable resistance in real-time we always kept the intensity of the electric field between the following values: 70-74 μA for *Amoeba proteus*; 70-80 μA for *Metamoeba leningradensis* and 68-75 μA for *Amoeba borokensis*.

The power supply was turned-off after 30 minutes of cell exposure. In the experiments in which a chemotactic gradient was not used the electrophoresis block utilized had never been in contact with any chemotactic substance.

A 5 minute galvanotaxis test was performed prior to any experiment that required the use of galvanotaxis. It is highly advisable to perform this check, as during experimentation we identified some cell populations that would not respond to galvanotaxis or even responded in an inverted manner, following to the anode.

### Cell induction

Only when all the cells in every experiment were attached to the glass surface, the laminar flux was established and the peptide nFMLP introduced in the left well of the second electrophoresis block. One minute later, the electric power supply was turned on. The process took 30 minutes and, after that, the power supply was turned off and only the cells that had moved towards the anode were rescued and placed in a Petri dish with clean Chalkley simplified medium for 10 minutes prior to future experiments. Using the ranges of electricity other than those specified in the previous section will give abnormal results as, for example, the ones depicted in Fig. S4. The electric field’s intensity values in that experiment ranged between 83 and 90 μA.

### Peptide gradient (chemotaxis)

After establishing the laminar flux, 750 μl of 2×10^−4^ M nFMLP (#F3506, Sigma-Aldrich) was added. The final concentration of the peptide solution was 2×10^−6^M. Immediately after introducing the peptide it was carefully mixed in the medium contained in the left well using a pipette until the amoebae started to move. Cell movements were recorded for 30 minutes.

### Peptide gradient calculation

The nFMLP peptide gradient was confirmed by measuring its concentration in the middle of the experimental chamber. 4μM fluorescein-tagged peptide (#F1314, Invitrogen) was loaded in the left side of the set-up. The set-up was prepared as usual but leaving a small opening (the size of the tip of a 50-200 μL micropipette) between the sliding cover glasses and the central glass piece. This opening allowed to get 60 μL samples from the central part of the laminar flux in the chamber at 0, 2, 5, 10, 15, 20 and 30 minutes following the establishment of the laminar flux. The peptide concentration was calculated extrapolating the values from a standard curve in which the concentration of the fluorescein-tagged peptide was known (Fig. S5). All the measurements were performed twice and the experiment was repeated three times. The fluorescence was measured in 96 well glass bottom black plates (P96-1.5H-N, In Vitro Scientific) employing a SynergyHTX plate reader (Biotek) at Excitation/Emission wavelengths of 460/528 following standard laboratory techniques as described by Green and Sambrook^38^.

### Track recording and digitizing

Cell trajectories were recorded using a digital camera linked to a SM-2T stereomicroscope. Images were taken every 10 seconds for a minimum of 30 minutes (180 frames). Only the first 30 minutes were quantified. As suggested by Hilsenbeck et al^39^, we did a manual tracking using the TrackMate software in ImageJ (http://fiji.sc/TrackMate)^40^, given the inaccuracies of automated tracking softwares^39^. Each track corresponds to the movements of an individual amoeba.

### Statistical significance

To estimate the significance of our results, we studied first if the distribution of cosines of angles came from a normal distribution, by applying the Kolmogorov-Smirnov test for single samples. Since the normality was rejected, the groups of cosines were compared by non-parametric tests, for groups by Kruskal-Wallis test, and pairs by the Wilcoxon rank-sum test, and therefore, the results were depicted as median/IQ instead of as mean±SD. Besides the p-values, we have reported the χ^2^ statistics and the Z statistics.

## Acknowledgments

We would like to thank Florentino Onandía Yague for valuable advice related to the experimental setup, and Laura Pérez Gómez for assistance in designing Fig. S1. We also express our gratitude to Dr. Alexey Smirnov (St. Petersburg University, Russia) for valued help with amoeba imaging. This work was supported by a grant of the University of Basque Country (UPV/EHU), GIU17/066, the Basque Government grant IT974-16, by the UPV/EHU and Basque Center of Applied Mathematics, grant US18/21, and by the Israel Science Foundation (536/19).

## Author contributions

JC-P, CB, and MF: performed the experiments; CB, MF and IMDF: design the setup; CB methodology with experimental glass chamber; MB and AG: cell cultures and cellular behavior advice; JC-P and CB: performed the digitalization of trajectories; IM: performed the quantitative studies; SK: performed the gradient analysis; SK, MDB and LM: main funding; MDB: laboratory facilities; SK, LM, MDB, APS, GPY, JIL and IM: analysis and design of the research mapping; all authors wrote the manuscript and agreed with its submission; IMDF: conceived, designed and directed the investigation.

---

**Figure S1.**
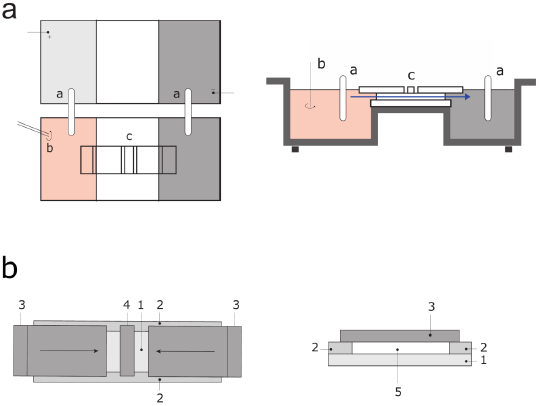
Experimental set-up. A illustrates the top and lateral views of the experimental system (two standard electrophoresis blocks), in red, the anode where the peptide is initially dissolved, in grey the cathode, a: agar bridges b: mixing pipette c: experimental glass structure; arrow illustrating the flow of the laminar flux. B shows the glass chamber where the cells are placed. 1: standard glass slide; 2: longitudinal cover glasses fixed to the glass slide; 3: sliding cover glasses; 4: central cover glass under which the cells are initially placed; 5: experimental chamber where a laminar flux is created and the cells can to migrate.

**Figure S2.**
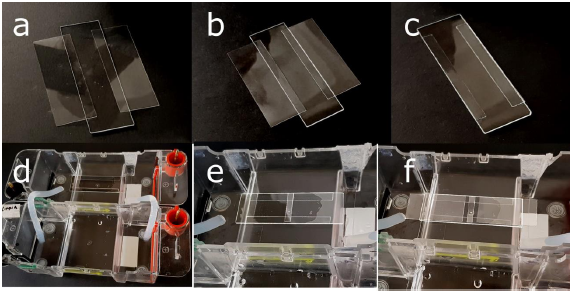
Experimental chamber. The specific set-up consists of two standard electrophoresis blocks, two agar bridges and a structure made from a standard glass slide and covers (the experimental chamber) Fig. S2d-f. The glass structure is composed of one slide and two covers (Fig. S2a). The two covers are fixed to the glass slide with silicone (Fig. S2b), and then they are trimmed with a methacrylate ruler (Fig. S2c). This modified slide is placed in the central platform of the second electrophoresis block (Fig. S2d). To avoid that the medium goes across the modified slide, we placed an oil drop under, it in the central platform of the block of electrophoresis. Finally, a central piece of cover glass about 3×24×0.1 mm (Fig. S2e) and two sliding lateral cover pieces are placed on the modified slide (Fig. S2f). When the sliding cover pieces are moved closing the central part (see Fig. S1) an inner laminar flux is generated in the chamber and when they are open, cell placement and rescue are possible under the central piece of coverslips.

**Figure S3.**
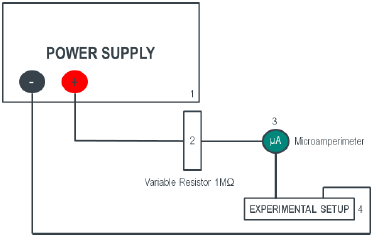
Supplementary electric device. The goal of these devices is to regulate and measure the electrical intensity that circulates through the system. 1, Standard Electrophoresis power supply, set at constant voltage. 2, 1MΩ linear variable resistor that regulates the current of the system. 3, a microamperimeter connected in series to the system, to measure the current intensity. 4, experimental setup described in Fig. 1 and Fig. S1.

**Figure S4.**
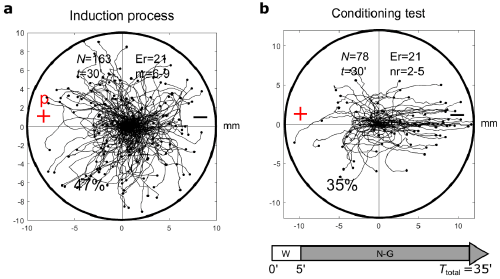
Conditioning process in *Amoeba proteus* using non-optimal electric field intensity. **a** Under simultaneous galvanotaxis and chemotaxis (induction process), 47% of the cells moved towards the anode-peptide (induced cells). These results agree with those obtained when using optimal intensity values for *Amoeba proteus* (70-74 μA, see Fig. 2). **b** After the induction process, the cells were placed in Chalkley’s medium without any stimulus for 5 min, and then they were exposed to galvanotaxis for 30 min using non-optimal electric field intensity values (83-90 μA). 35% of the induced cells presented lasting directionality towards the anode (where the chemotactic peptide was absent) compared to 75% when using optimal intensity values for *Amoeba proteus* (see Fig. 2). However, despite using non-optimal calibration, the cosine values obtained from the conditioned test were significantly different to those from the galvanotaxis (*p* − *value* = 10^−5^), indicating the emergence of a new behaviour in the migration patterns. “N” total number of cells, “Er” experimental replicates, “nr” number of cells per replicate, “t” time of galvanotaxis or chemotaxis, “+” anode, “−” cathode. Both the x and y axis show the distance in mm, and the initial location of each cell has been placed at the center of the diagram.

**Figure S5.**
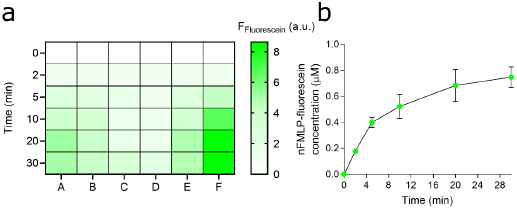
Fluorescein-tagged peptide concentration in the middle part of the laminar chamber flux. Fluorescein-tagged peptide concentration in the middle part of the laminar chamber flux as determined with a microplate reader. **a**, heat map presenting individual fluorescence measurements at different times. **b**, average levels of nFMLP-fluorescein concentration as time-function. The data represent the Mean±SEM of 6 measurements (A-F, taken at 0, 2, 5, 10, 20, and 30 min). The peptide concentration in the middle of the glass experimental chamber (where the amoebae are placed) increases immediately following the flow establishment (within 2 minutes the concentration rises from zero to approximately 0.2 μM) and this concentration increases further (to 0.6 μM) for at least 30 minutes.

## References

1 Hawkins, R. D. & Byrne, J. H. Associative learning in invertebrates. Cold Spring Harb. Perspect. Biol. 7, a021709 (2015).

2 De la Fuente, I. M. et al. Evidence of conditioned behavior in amoebae. Nat. Commun. 10, 3690 (2019).

3 Yudin, A.L. Animal Species for Developmental Studies, 1–11. (Springer US, Boston, 1990).

4 Jeon, K. W. The large, free-living amoebae: wonderful cells for biological studies. J. Eukaryot. Microbiol. 42, 1–7 (1995).

5 Goodkov, A., Yudin, A. & Podlipaeva, Y.I. Collection of the proteus-type amoebae at the Institute of Cytology, Russian Academy of Sciences. I. History, goals and research fields. Protistology 8, 71–75 (2014).

6 Berdieva, M., Demin, S. & Goodkov, A. Amoeba proteus and ploidy cycles: From simple model to complex issues. Protistology 13, 166–173 (2019).

7 Grebecki, A. Behaviour of Amoeba proteus exposed to light-shade difference. Protistologica 16, 103–113 (1980).

8 Korohoda, W., Golda, J., Sroka, J. & Wojnarowicz, A. Chemotaxis of Amoeba proteus in the developing pH gradient within a pocket-like chamber studied with the computer assisted method. Cell Motil. Cytoskelet. 38, 38–53 (1997).

9 Korohoda, W., Mycielska, M., Janda, E. & Madeja, Z. Immediate and long‐term galvanotactic responses of Amoeba proteus to dc electric fields. Cell. Motil. Cytoskelet. 45, 10–26 (2000).

10 Smirnov, A. Encyclopedia of Microbiology, 558–577. (Elsevier, Oxford, 2009).

11 Lugini, L. et al. Cannibalism of live lymphocytes by human metastatic but not primary melanoma cells. Cancer Res. 66, 3629–3638, (2006).

12 Fais, S. & Fauvarque, M.-O. TM9 and cannibalism: how to learn more about cancer by studying amoebae and invertebrates. Trends Mol. Med. 18, 4–5 (2012).

13 Demin, S. Y., Berdieva, M. & Goodkov, A. Cyclic polyploidy in obligate agamic amoebae. Cell Tissue Biol. 13, 242–246 (2019).

14 Erenpreisa, J. et al. Stress-induced polyploidy shifts somatic cells towards a protumourogenic unicellular gene transcription network. Cancer Hypotheses 1, 1–20 (2018).

15 Bray, D. Cell Movements: From Molecules to Motility. (Garland Science, New York, 2000).

16 Prusch, R. D. & Britton, J. C. Peptide stimulation of phagocytosis in Amoeba proteus. Cell Tissue Res. 250, 589–593 (1987).

17 De la Fuente, I.M. et al. The nucleus does not significantly affect the migratory trajectories of amoeba in two-dimensional environments. Sci. Rep. 9, 1–15 (2019).

18 De la Fuente, I. M. Metabolic Dissipative Structures in Systems Biology of Metabolic and Signaling Networks: Energy, Mass and Information Transfer, 179–212. (Springer Berlin Heidelberg, 2014).

19 De la Fuente, I. M., Cortes, J. M., Pelta, D. A. & Veguillas, J. Attractor metabolic networks. PLoS ONE 8, e58284 (2013).

20 De la Fuente, I. M. Elements of the cellular metabolic structure. Front. Mol. Biosc. 2, 16 (2015).

21 De la Fuente, I. M. & López, J. I. Cell motility and cancer. Cancers 12, 2177 (2020).

22 Finger, S. Origins of Neuroscience: A History of Explorations into Brain Function. (Oxford Univ. Press, Oxford, 1994). Revised, 2001.

23 Pavlov, I. P. Conditioned reflexes: an investigation of the physiological activity of the cerebral cortex. (Oxford Univ. Press, Oxford, 1927).

24 Dunigan, D. D. et al. Chloroviruses Lure Hosts through Long-Distance Chemical Signaling. J. Virol. 93, e01688–18 (2019).

25 Rescorla, R. A. Pavlovian conditioning and its proper control procedures. Psychol. Rev. 74, 71–80 (1967).

26 Gibbs, J. W. Elementary Principles in Statistical Mechanics Developed with Especial Reference to the Rational Foundation of Thermodynamics. (Charles Scribner’s Sons, New York, 1902).

27 Einstein, A. Zum gegenwartigen Stand des Strahlungsproblesm. Phys. Zeits. 10, 185–193 (1909).

28 Tagkopoulos, I., Liu, Y.-C. & Tavazoie, S. Predictive behavior within microbial genetic networks. Science 320, 1313–1317 (2008).

29 Armus, H. L., Montgomery, A. R. & Jellison, J. L. Discrimination learning in paramecia (P. caudatum). Psychol. Rec. 56, 489–498 (2006).

30 Boisseau, R. P., Vogel, D. & Dussutour, A. Habituation in non-neural organisms: evidence from slime moulds. Proc. R. Soc. B 283 (2016).

31 Shirakawa, T., Gunji, Y. P. & Miyake, Y. An associative learning experiment using the plasmodium of Physarum polycephalum. Nano Commun. Netw. 2, 99–105 (2011).

32 Sanchez-Andres, J. V., Pomares, R. & Malaisse, W. J. Adaptive short-term associative conditioning in the pancreatic β-cell. Physiol. Rep. 8, e14403 (2020).

33 Moore, B. R. The evolution of learning. Biological Reviews. 79:301–335 (2004).

34 Stuelten, C. H., Parent, C. A. & Montell, D. J. Cell motility in cancer invasion and metastasis: insights from simple model organisms. Nat. Rev. Cancer 18, 296–312 (2018).

35 Mezu-Ndubuisi, O.J., Maheshwari, A. The role of integrins in inflammation and angiogenesis. Pediatr. Res. (2020).

36 Li, L. et al. Long noncoding RNA SNHG7 accelerates proliferation, migration and invasion of non-small cell lung cancer cells by suppressing miR-181a-5p through AKT/mTOR signaling pathway. Cancer Manag. Res. 12, 8303–8312 (2020).

37 Prescott, D.M. & Carrier, R.F. Experimental procedures and cultural methods for Euplotes eurystomus and Amoeba proteus. Methods Cell Biol. 1, 85–95 (1964).

38 Green, M.R. & Sambrook, J. Molecular Cloning, A Laboratory Manual, Fourth Edition (Cold Spring Harbor Laboratory Press, New York, 2012).

39 Hilsenbeck, O. et al. Software tools for single-cell tracking and quantification of cellular and molecular properties. Nat. Biotechnol. 34, 703–706 (2016).

40 Tinevez, J. Y., et al. TrackMate: An open and extensible platform for single-particle tracking. Methods. 115, 80–90 (2017).

